# Structural Analysis of Phosphorylation Proteoforms in a Dynamic Heterogeneous System Using Flash Oxidation Coupled In-Line with Ion Exchange Chromatography

**DOI:** 10.1101/2022.10.04.510855

**Authors:** Zhi Cheng, Sandeep K. Misra, Anter Shami, Joshua S. Sharp

## Abstract

Protein post-translational modifications (PTMs) are key modulators of protein structure and function that often change in a dynamic fashion in response to cellular stimuli. Dynamic post-translational modifications are very challenging to structurally characterize using modern techniques, including covalent labeling methods, due to the presence of multiple proteoforms and conformers together in solution. Here, we have coupled ion exchange HPLC with a flash oxidation system (IEX LC-FOX) to successfully elucidate structural changes among three phosphoproteoforms of ovalbumin (OVA) during dephosphorylation with alkaline phosphatase (AP). Real-time dosimetry indicates no difference in effective radical dose between peaks or across the peak, demonstrating both the lack of scavenging of the NaCl gradient and the lack of a concentration effect on radical dose between peaks of different intensities. The use of IEX LC-FOX allows us to structurally probe each phosphoproteoform as it elutes from the column, capturing structural data before the dynamics of the system reintroduce heterogeneity. We found significant differences in residue-level oxidation between the hydroxyl radical footprint of non-phosphorylated, mono-phosphorylated and di-phosphorylated ovalbumin. Not only were our data consistent with the previously reported stabilization of ovalbumin structure by phosphorylation, but local structural changes were also consistent with the measured order of dephosphorylation of Ser344 being removed first. These results demonstrate the utility of IEX LC-FOX for measuring the structural effects of PTMs, even in dynamic systems.

## INTRODUCTION

PTMs play an essential role in creating complexity within the proteome for diverse functions using only a limited number of genes ^1,2^. PTMs can influence almost all aspects of cell biology and pathogenesis by influencing the protein structure, stability, activity, cellular localization, or substrate specificities ^3-5^. Some PTMs reversibly control protein functions, such as cell signaling, by being dynamically added and removed ^6^. Therefore, understanding how PTMs alter a protein’s structure and function in a dynamic system is crucial for disease prevention and drug discovery. Our knowledge of existing PTMs is wide and still expanding, but one of the most widely studied PTMs is protein phosphorylation. Protein phosphorylation plays an essential role in mammalian biology. About 14,000 proteins encoded by the human genome are shown to be phosphorylated. Phosphorylation is extremely important in most cellular processes such as protein synthesis, cell division, signal transduction, cell growth, development, and aging^7^. Mutation or defects in phosphorylation regulatory mechanisms can lead to abnormal activation of the kinase signaling pathway, potentially resulting in serious issues in cellular physiology such as cancer, developmental disorder, and degenerative diseases^8^. However, the structural effects of these PTMs are often extremely difficult to analyze because they exist in a dynamic system that changes protein PTM status and conformation over time, resulting in a mixture of proteoforms, each with its own conformation. Hydroxyl radical protein footprinting (HRPF) is an MS-based method for measuring protein topography, based on generating hydroxyl radicals *in situ* and allowing them to modify the surface amino acids of the protein. The extent of modification at a given amino acid depends on the side chain’s solvent accessibility as well as its inherent reactivity towards hydroxyl radicals ^9^. HRPF coupled with LC-MS has been used rapidly in protein structural studies in recent decades, such as identifying higher-order protein structure, protein-protein interactions, and protein-ligand interactions ^10-22^. However, in current HRPF methods, systems that contain heterogeneous mixtures of conformers due to structural dynamics or the presence of multiple proteoforms result in data that reflects the weighted average of all conformations in solution, frustrating analysis of these systems^23^. The structure of any single conformer in the heterogeneous mixture remains unknown.

Our group recently published a study in which we developed HPLC size exclusion chromatography (SEC) coupled with a traditional excimer laser-based fast photochemical oxidation of proteins (FPOP) system to successfully analyze the monoclonal antibody adalimumab in its native and aggregated conformations^23^. However, this method does not allow us to analyze the structural effects of most PTMs due to the relatively poor resolution of SEC. Additionally, HPLC coupled with the traditional FPOP setup was very bulky and needed to deal with significant laser hazards and hazardous fluorine gas. SEC uses an isocratic gradient, which provides a constant radical scavenging background; it remains unclear if the changing buffer compositions commonly found in higher resolution gradient-based protein chromatography would confound inline HRPF by introducing a changing radical scavenging background. Here, we tested the recently introduced FOX photolysis system^24^ coupled with HPLC weak anion exchange chromatography (WAX) using a sodium chloride gradient to separate and label each ovalbumin (OVA) phosphoproteoform in a real-time dynamic system of alkaline phosphatase acting on OVA^25^. This system allows us to determine if IEX LC-FOX is capable of measuring structural differences in a dynamically-changing mixture of phosphoproteoforms.

## EXPERIMENTAL SECTION

### Materials

Ovalbumin (OVA), alkaline phosphatase (AP), equine myoglobin, [Glu]1-fibrinopeptide B (GluB), iodoacetamide (IAA), catalase, methionine amide, calcium chloride (CaCl_2_), sodium chloride (NaCl) and formic acid (FA) were purchased from Sigma-Aldrich Crop. (St. Louis, MO). Adenine, dithiothreitol (DTT), LC/MS grade acetonitrile and water, mono- and di-basic sodium phosphate, and Tris (hydroxymethyl) aminomethane were purchased from Fisher Scientific (Fair Lawn, NJ). Hydrogen peroxide (30%) was purchased from JT Baker (Phillipsburg, NJ). Fused silica capillary was purchased from Digi-Key (Thief River Falls, MN). Sequencing grade modified trypsin was purchased from Promega (Madison, WI). PNGase F was purchased from New England BioLabs (Ipswich, MA).

### LC-FOX Instrument Setup

The effluent line leaving the UV detector of a Dionex Ultimate 3000 HPLC (Thermo Fisher, CA) system was connected to a microtee connector (IDEX, IL). The microtee connector used 250 μm fused silica capillaries for both pathways: a short eluant line leading to a second microtee, and a resistor line going to waste, creating a 1:20 split to reduce the flow rate (FR) from 200 μL/min leaving the HPLC UV detector to 10 μL/min entering the second microtee connector. In the second microtee, a make-up flow of 5 μL/min of 300 mM hydrogen peroxide was mixed with the eluant at a 1:2 ratio, for a final concentration of 100 mM hydrogen peroxide entering the photolysis cell at 15 μL/min. Analytes were oxidized and passed to an inline dosimeter to monitor the effective hydroxyl radical dose as previously described^24,26^. Finally, each analyte was collected separately in the quenching solution in the product collector (**Figure 1**).

**Figure 1.**
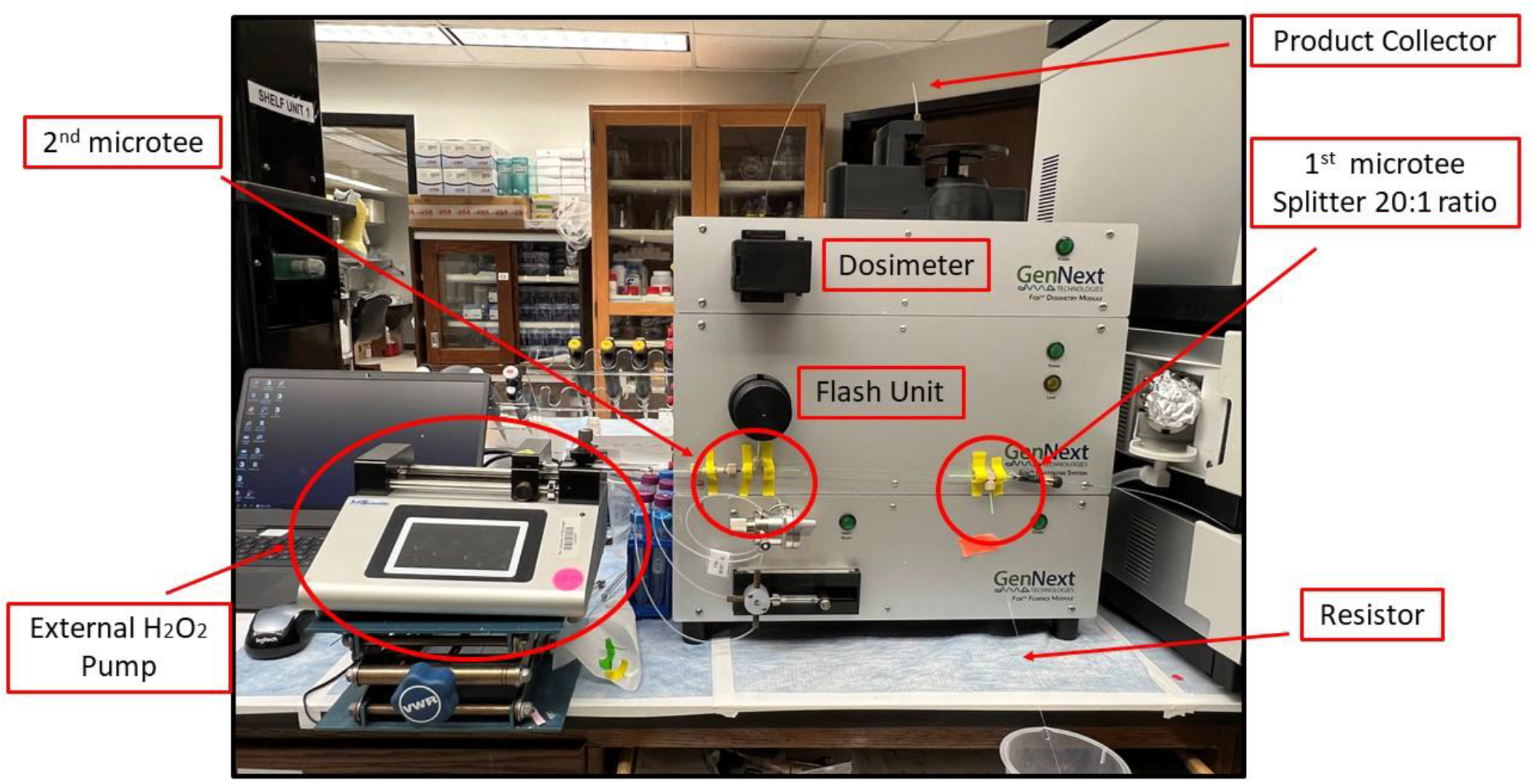
HPLC-FOX instrumentation setup. The HPLC system is to the right of the frame. The fluidics module of the Fox Photolysis System is bypassed, with the second microtee output leading directly to the flash photolysis cell.

### Measurement of the radical scavenging capacity of ion exchange salt gradient

FOX samples contained 50 mM sodium phosphate (pH 7.4), 2 mM adenine, 5 μM myoglobin, 5 μM GluB, 100 mM hydrogen peroxide, and various concentrations of NaCl (0, 0.1, 0.3, 0.6, 1 M). Samples were prepared in triplicate and irradiated by FOX with a flash voltage of 600 V at 2 Hz (20% exclusion volume). Real-time changes in adenine absorption at 265 nm (ΔAbs265) were measured to measure the effective radical dose in the presence of each salt concentration^26-28^. A decrease in ΔAbs265 indicate increased hydroxyl radical scavenging. 12 μL of each sample was collected into 25 μL of quenching solution containing 0.3 mg/mL catalase and 35 mM methionine amide to quench the excess H_2_O_2_ and prevent secondary oxidation. Quenched myoglobin samples were digested with trypsin and analyzed in LC-MS under the method previously publised^29^.

### LC-FOX with OVA Phosphoproteoform Analysis

100 μL of 50 μg/μL OVA was treated with 111 units of AP at room temperature. Triplicate OVA+AP samples (1 mg) after a reaction time between 40-100 min, which were found to have intermediate levels of dephosphorylation, were loaded onto a PolyLC (Columbia, MD) WAX column (100 × 2.1 mm, 5 µm, 1000 Å) with a flow rate of 200 μL/min. Solvent A was 12.75 mM Tris and solvent B was 12.75 mM Tris with 1 M NaCl (pH 6.9 for both solvents). The gradient consisted of 100% A held for 2 min, 2-50% solvent B over 21 min, 50-80% B over 1 min, held at 80% B for 3 min, returned to 100% A over 1 min and held at 100% A for 4 min. HPLC UV detection wavelength was set at 280 nm. Each peak of the OVA proteoform was irradiated by the Fox System with a flash voltage of 800 V at 2 Hz for 36 seconds around the apex. The FOX inline dosimeter measured the change in Tris absorption at 265 nm to ensure that comparable hydroxyl radicals were generated among the replicates^30^. 9 μL of each oxidized proteoform was collected into 25 µL of quenching solution containing 0.3 mg/mL catalase and 35 mM methionine amide to quench the excess H_2_O_2_ and prevent secondary oxidation.

Repeated LC-MS/MS analysis of the AP-treated samples indicated a complete loss of phosphorylation at Ser344 for all phosphoproteoforms resolved by WAX, and loss of ∼65% of phosphorylation at Ser68 even for the diphosphorylated proteoforms (data not shown). These results indicated that AP dephosphorylation continued after WAX separation in an ongoing dynamic reaction, confounding our ability to quantify phosphorylation directly from the AP-treated samples. Therefore, we examined phosphorylation on naturally-occurring phosphoproteoforms for untreated OVA, using WAX retention time to verify that the phosphoproteoform collected was the same as observed from AP treatment. To identify the three proteoforms observed by WAX, triplicate OVA samples were prepared in water and separated by WAX as described above. Four peaks were observed, three of which matched with peaks observed after AP treatment (with the one new peak eluting between the monophosphorylated and dephosphorylated proteoforms showing ∼60% phosphorylation at both sites, data not shown). Each peak was collected in a 30 second fraction centered at the peak apex, digested with trypsin, and analyzed by LC-MS (described in the LC-MS analysis section). The percentage of the phosphorylation in each peak corresponding to the peaks observed in AP-treated samples was calculated based on two tryptic peptides 59-84 and 340-359, which included Ser-68 and Ser-344. From the LC-MS result, two different charge states were found for each peptide, and the signal from the two charge states was summed. The percentage phosphorylation was calculated manually with Xcalibur V.3.1 using Equation 1, where *I* represents the integrated peak intensity of a phosphoform of one of the two peptides.

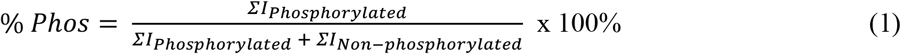

### LC-MS Analysis of Oxidized Samples

After LC-FOX, samples were diluted into 50 mM Tris containing 1 mM of CaCl_2_ (pH 8.0) and 10 mM DTT. Samples were heated at 95 °C for 30 min then cooled down to room temperature. 2 μL of 500 mM IAA was added to each sample and reacted in the dark for 30 min. 1 μL of 500 mM DTT was added to each sample to quench IAA. For protein digestion, a 1:20 ratio of trypsin/OVA was added to the samples and incubated at 37 °C overnight with sample mixing. Trypsin was inactivated by heating samples at 95 °C for 10 min. For protein deglycosylation, PNGase F was added at a 50 unit/μg protein ratio and incubated at 37 °C for an hour. 1 μL of 500 mM DTT and 0.1% formic acid were added to each sample before LC-MS analysis. Digested OVA was analyzed by an Orbitrap Fusion Tribrid mass spectrometer coupled with a Dionex Ultimate 3000 system nanoLC system (Thermo Fisher, CA). OVA samples were loaded onto a trap column (Acclaim PepMap C18 5 um, 0.3 × 5 mm) and eluted onto a nano C18 column (Acclaim 3 µm, 0.075 × 150 mm) at a flow rate of 0.3 µL/min. Solvent A was 0.1% formic acid in water, and solvent B was 0.1% formic acid in acetonitrile. The gradient consists of 2% solvent B held for 5 min, 2-10% B over 5 min, 10-50% B over 27 min, 50-95% B over 2 min and held for 5 min, returned to 2% B over 3 min and held for 9 min. ESI was carried out in positive ion mode, with a nominal orbitrap resolution of 60000 and an RF lens setting of 60. Data-dependent mode MS/MS was used, with both CID and EThcD applied to each selected precursor. CID intensity threshold was set at 5.3e3, and CID collision energy was set at 30% with a rapid ion trap scan. EThcD intensity threshold was set at 2.5e4, and supplementary activation collision energy set at 6% with an orbitrap nominal resolution of 30000. Byonic (v4.4.1, Protein Metrics, San Carlos, CA) was used to identify the peptide and determine the protein sequence coverage. Average oxidation event per peptide and residue were calculated manually with the help of Xcalibur, V.3.1 as previously published ^29,31^. For residue level analysis, the *m/z* and retention times of OVA peptides and peptide oxidation products were noted from the initial DDA run and placed in an inclusion list (Supplemental Table 1). Targeted mass fragmentation was by CID with a collision energy of 30%, and the detector type was an ion trap. Sites of oxidation were assigned manually from MS/MS spectra for each oxidized peak, and peak area was used to calculate residue level oxidation ^32^. The mass spectrometry proteomics data have been deposited to the ProteomeXchange Consortium via the PRIDE partner repository ^33^ with the dataset identifier PXD037050.

### Size Exclusion Chromatography coupled with Multi-Angle Light Scatting (SEC-MALS) Analysis

AP treated and untreated OVA samples were prepared under the same conditions described in the LC-FOX section above. SEC-MALS were performed with a Dionex Ultimate 3000 system (Thermo Fisher, CA) coupled with a miniDAWN Treos multi-angle light scattering detector (Wyatt, CA). 1 mg of OVA sample was loaded onto a size-exclusion column (ACQUITY UPLC Protein BEH 150 mm, 125Å, 1.7 μm, Waters, Milford, MA). 100 mM of sodium phosphate at pH 7.4 was used as the mobile phase, and the separation was under an isocratic gradient at a flow rate of 0.2 mL/min. The UV wavelength was set at 280 nm. Data from the SEC-MALS were analyzed using ASTRA 7.1.3.

### Molecular dynamics simulations

The initial coordinates of non-phosphorylated ovalbumin were taken from the Protein Data Bank (PDB ID: 1OVA)^34^. The input files for MD simulation were prepared using CHARMM–GUI glycan modeler^35^ on CHARMM–web^36^. The missing residues were modeled using the GalalyxFill^37^ option in the CHARMM– web. Residues Ser68 and Ser344 were phosphorylated using CHARMM–web. A non-phosphorylated and two mono-phosphorylated proteoforms were generated. A rectangular TIP3P water box was added, extending 10 Å in each dimension from the edges of the protein. Cell-like isotonic conditions were maintained by adding 0.15 M NaCl ions (placed using the Monte-Carlo approach) to each system. Both protein and glycans were treated with the CHARMM36m force-field. Both systems were then subjected to multistep equilibration^38^ followed by 1 μs MD simulation at constant NPT using T = 300 K; temperature scaling by Langevin dynamics; 2 ps collision frequency; pressure relaxation every 1.2 ps; SHAKE constraints; non-bonded interaction cutoff of 9 Å and an integration time step of 2 fs. The *cuda* version of *pmemd* in Amber20^39^ was used to perform equilibration and the production MD runs. Atomic coordinates were saved every 1 ns and were subsequently analyzed by *VMD*^*40*^.

## RESULT AND DISCUSSION

### NaCl Gradients Do Not Scavenge Hydroxyl Radicals

The ability of NaCl to act as a hydroxyl radical scavenger in the FOX system was tested to determine if radical compensation must be performed^16^. In order to ensure that samples received the same effective hydroxyl radical dose, regardless of the position in the gradient, we measured the hydroxyl radical scavenging capabilities of NaCl in the concentration range from 0-1 M. Both inline dosimetry and peptide and protein oxidation were used to evaluate the effect of NaCl concentration on effective hydroxyl radical dose.

Five different concentrations of NaCl (0, 0.1, 0.3, 0.6, and 1 M) were added to myoglobin samples and FOX was performed. Inline adenine radical dosimetry showed no significant scavenging effect by any NaCl concentration in the range of 0-1 M (**Figure 2A**). Examination of the oxidation of both myoglobin and GluB by LC-MS data confirmed there was no significant radical scavenging by NaCl at any concentration tested (**Figure 2B**). Based on these results, no radical compensation was performed to correct for NaCl gradient scavenging in OVA experiments.

**Figure 2.**
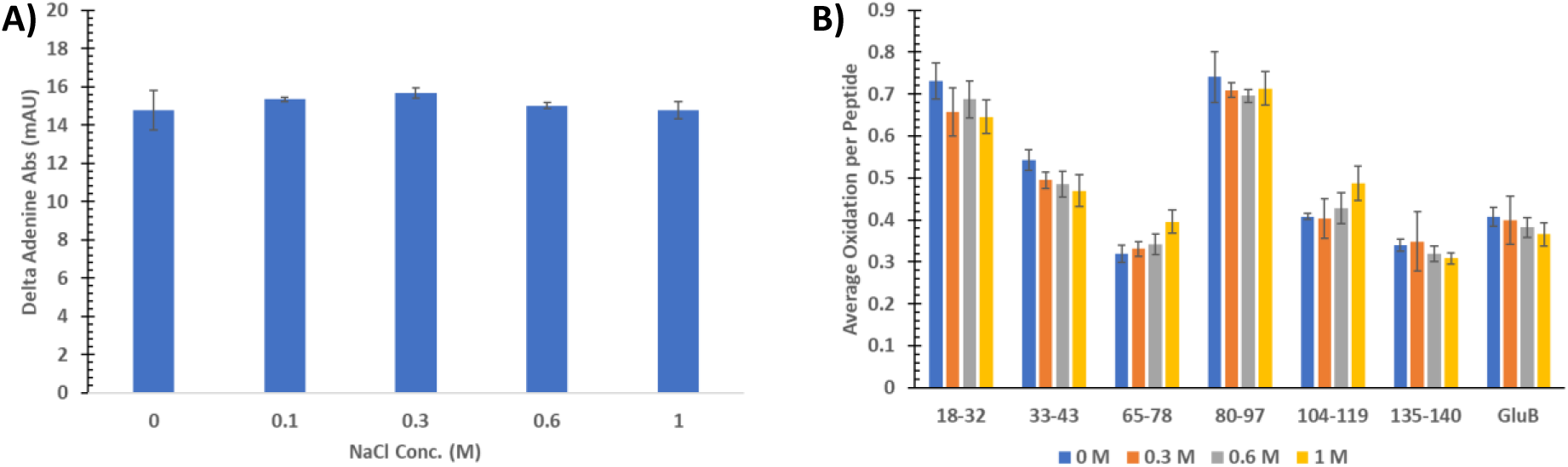
Measurement of hydroxyl radical scavenging by NaCl at concentrations up to 1M. **A)** FOX inline dosimetry measurements for triplicate myoglobin samples at different concentrations of NaCl. **B)** LC-MS analysis for FOX oxidation of myoglobin and GluB in the presence of different concentrations of NaCl. Error bars represent one standard deviation.

### LC-FOX for OVA Phosphorylated Proteoforms Analysis in the Dynamic System

To test the ability of IEX LC-FOX to structurally analyze phosphoproteoforms in a dynamic system, alkaline phosphatase (AP) was added to the OVA. AP enzymatically hydrolyzes phosphate groups from the proteins, and the ongoing reaction was analyzed as a model dynamic system for OVA proteoforms analysis. By using the IEX chromatography, we were able to separate three phosphoproteoforms of OVA that changed in abundance as the reaction proceeded (**Figure 3A**). We selected 36 seconds around the apex of each phosphoproteoform peak and immediately irradiated it with the inline FOX UV system as it eluted from the column using the modified system shown in **Figure 1**. All the proteoforms were collected individually into the quenching solutions between the OVA+AP reaction time between 40-100 min range.

**Figure 3.**
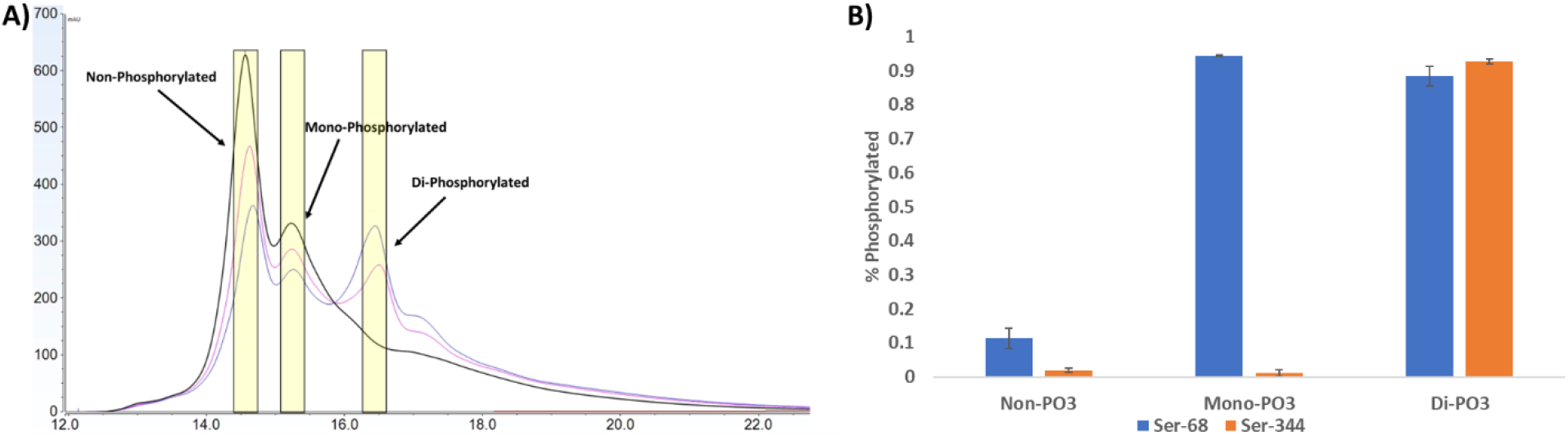
HPLC and LC-MS analysis of OVA-AP reaction. **A)** WAX separation of OVA-AP reaction after (blue) 40 minutes, (magenta) 130 minutes or (black) 48 hours. Each peak represents a different phosphoproteoform of OVA in the dynamic system. The regions of each peak subjected to FOX are highlighted in yellow. **B)** LC-MS results identify the three peaks from the IEX separation as non-, mono-, and di-phosphorylated proteoforms, with the monophosphorylated form being almost solely phosphoSer68.

Bottom-up LC-MS identification of the corresponding peaks from untreated OVA confirmed that the first peak was non-phosphorylated (non-PO3) OVA, the middle peak was mono-phosphorylated (mono-PO3) OVA, and the third was di-phosphorylated (di-PO3) OVA. More specifically, the mono-PO_3_ OVA was almost solely phosphoSer-68 (**Figure 3B**). To determine if the AP reaction would alter the OVA monomer-dimer equilibrium^25^, the OVA monomer and dimer ratio was measured by SEC-MALS. The UV trace from HPLC indicated that the OVA monomer(s) and dimer ratio was at the same level in all samples with ∼13% dimer and ∼87% monomer(s). **Figure S1A** showed regardless of the AP treatment or the AP reaction time, the ratio of the OVA monomer and dimer was consistent. According to these results, we can conclude that both the treatment of the AP and the time of the AP reaction did not cause the OVA dimerization. We also observed with AP acting on OVA, both UV **(Figure S1A)** and MALS **(Figure S1B)** traces showed additional monomer peak from the OVA monomer increased proportionally with AP reaction time. We think the different OVA monomer peaks indicated the loss of the phosphoryl group from OVA during the AP dynamic system to causing the structure to undergo a conformational change. Therefore, different sizes of the OVA monomers were able to be separated from the SEC system.

### Molecular Dynamics Simulations of Local Impacts of Phosphorylation

Previous reports demonstrated that phosphorylation of OVA generally stabilizes the higher order structure^41^. This role of phosphorylation in OVA dynamics leads us to hypothesize that dephosphorylation will generally lead to higher levels of oxidation in many regions of the protein with little indication of decreased oxidation in the dephosphorylated samples. However, such a hypothesis lacks specificity and is not ideal for validating our LC-FOX technique. Before we analyzed each proteoform of OVA by LC-FOX, we performed a 1 μs all-atom molecular dynamics simulation for each OVA phosphoproteoform to generate testable hypotheses regarding structural effects of phosphorylation. Non-phosphorylated, phosphoSer-68, and phosphoSer-344 OVA were simulated as described. The brief simulation should reliably capture local changes in structure and dynamics caused directly by phosphorylation, without attempting to accurately simulate more systemic protein stabilization effects that have been reported^42,43^. We observed one significant local structural difference between non-phosphorylated and phosphorylated Ser-344 (**Figure 4**). Ser344 is located on a loop. MD simulation showed that the phosphate group from phosphoSer344 coordinated with Lys189 in a neighboring loop, which stabilized the Ser344 loop. Without phosphorylation, this interaction did not occur and the loop that contains Ser344 was highly dynamic. We hypothesized that dephosphorylation should impact oxidation of amino acids on the Ser344 loop and/or show significant protection of the adjacent interacting loop and strands, consistent with the short MD simulation.

**Figure 4.**
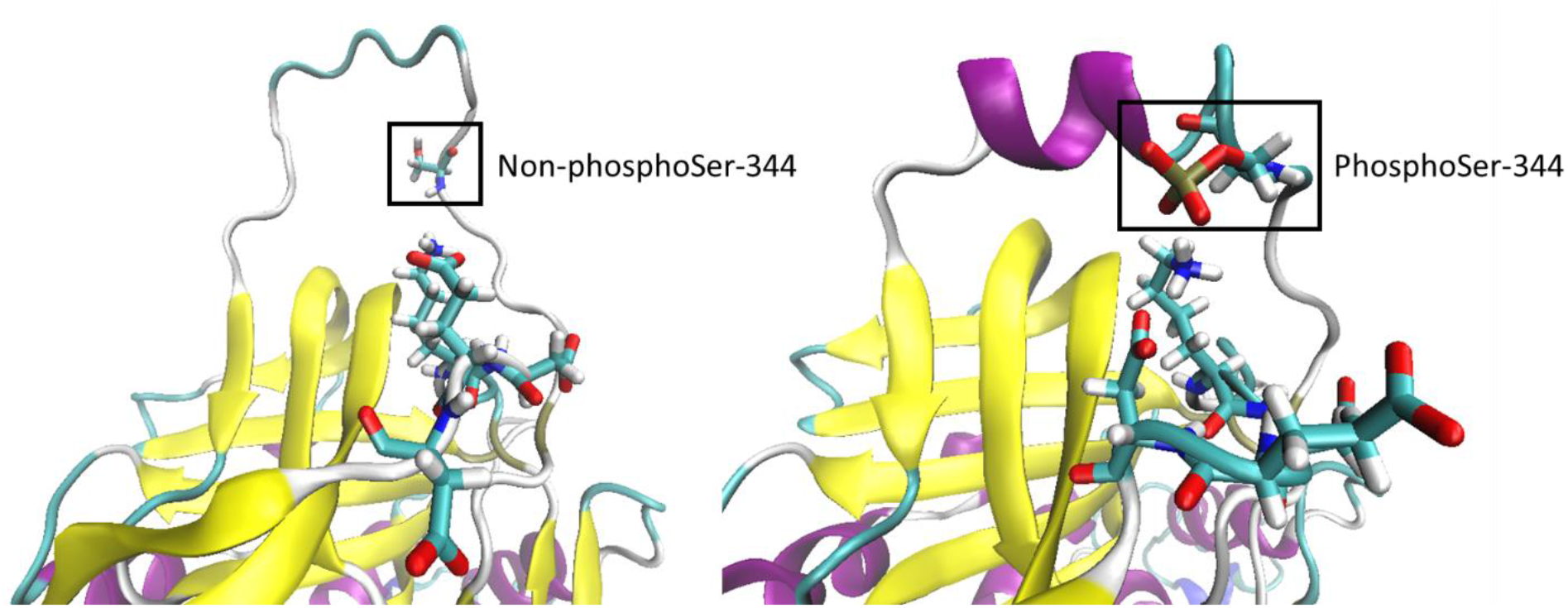
Molecular dynamics simulation results show a significant difference between non-phosphoSer-344 and phosphoSer-344. The phosphoSer-344 showed the phosphate group interacted with Lys189 to stabilize the loop; when Ser344 is dephosphorylated, the loop is unstructured.

### IEX LC-FOX analysis of OVA phosphoproteoforms at the peptide and residue level oxidation

LC-FOX samples were digested with trypsin and analyzed by LC-MS/MS. The MS results of peptide level oxidation calculation are shown in **Figure S2**. One-way ANOVA with a 95% confidence interval statistical test was applied for each peptide. Seventeen peptides showed oxidation, and ten of these peptides showed a significant decrease in oxidation (1-16, 20-46, 85-104, 105-123, 127-142, 143-158, 191-200, 201-219, 229-263, and 323-339) among at least one of the three OVA phosphoproteoforms compared to the fully dephosphorylated proteoform. These results are consistent with our general hypothesis that dephosphorylation should result in more oxidation at many regions of the protein, consistent with destabilization of the protein structure^41^.

Three different patterns of oxidation were observed in the peptides, reflecting the roles of phosphorylation in structure. The first pattern showed that significant protection occurred upon phosphorylation of Ser68 OVA, with no additional protection observed upon phosphorylation of Ser344 (20-46, 105-123, 127-142, 143-158, and 201-219). The second pattern showed no difference between nonphosphorylated and monophosphorylated forms, with protection only occurring upon phosphorylation of Ser344 in the dephosphorylated sample (85-104, 191-200, and 323-339). The third pattern showed protection upon phosphorylation of Ser68, which increased upon phosphorylation of Ser344 indicating additive protection from each phosphorylation. These results are in agreement with our general hypothesis, based on previously reported results that phosphorylation stabilizes OVA structure.

To test our specific hypotheses regarding the targeted role of phosphorylation of Ser344 on the local structure, we examined peptides in spatial vicinity of Ser344. Peptide 191-200 contains the amino acids from the neighboring loop that coordinates with the Ser-344 phosphate group as observed in the simulation experiment. This peptide showed significant protection only between non-phosphorylated and di-phosphorylated OVA, which is the only sample that contains phosphoSer-344. This result supports our MD-based hypothesis that phosphorylation of Ser-344 should lead to protection of the neighboring loop containing Lys189, and suggests that LC-FOX is generating data consistent with expectations for OVA phosphoproteoforms.

In order to obtain higher spatial resolution for our LC-FOX results, we performed residue-level analysis. Residue level data was obtained from MS data-dependent acquisition (DDA) with a targeted mass list generated from initial LC-MS runs (**Supplementary Table 1**). 15 of 27 amino acids from the residue level calculation showed significant decreases in oxidation level upon OVA phosphorylation (**Figure 5**). Nine residues or short regions (C30, S98, RY110-111, F134, W148, S151, V327, H328, and I344) showed significantly decreased oxidation in mono-phosphorylated compared to non-phosphorylated OVA but no significant difference between the mono-phosphorylated vs. di-phosphorylated OVA, which represented the first protection pattern that we described in peptide level analysis. Six residues (M8, D95, T136, M196, M210, and M211) showed significant protections in di-phosphorylated compared to both

**Figure 5.**
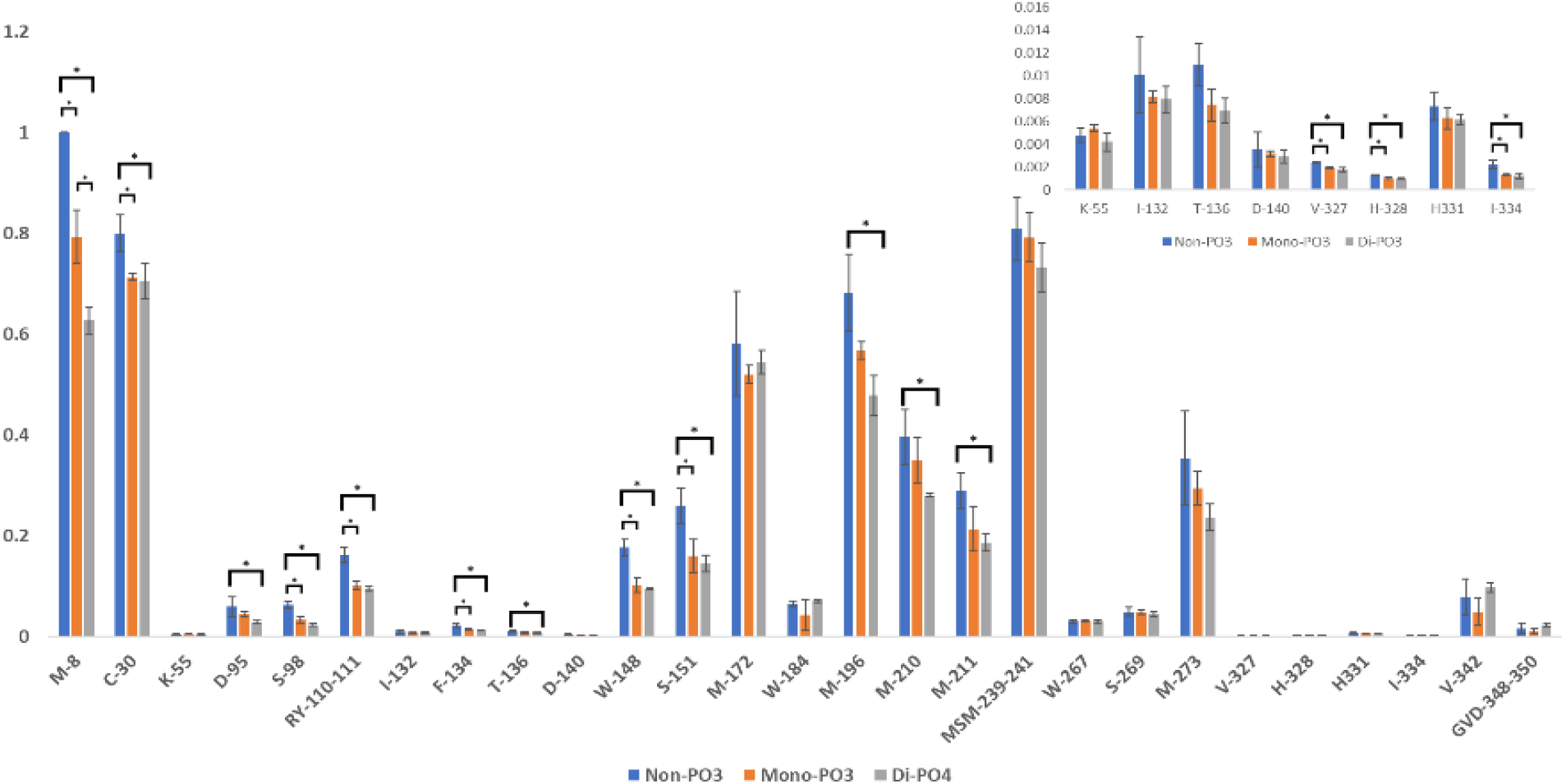
Residue level oxidation calculation of three OVA proteoforms from IEX LC-FOX. Asterisks mark data pairs with statistically significant differences, as indicated by the bracket. Significant difference was tested by one-way ANOVA followed by Tukey post-hoc analysis (α=0.05). non-phosphorylated and mono-phosphorylated OVA, indicating the protections were due to the additional Ser-344 phosphorylation, as described in data for peptide level oxidation analysis.

To contextualize the changes in these amino acids with the structure of OVA, we mapped the OVA LC-FOX data to the X-ray crystal structure of OVA at 1.95 Å^34^ using PyMol^44^ (**Figure 6**). Two comparison models were made based on the LC-MS result: non-PO3 vs. mono-PO3 and non-PO3 vs. di-PO3. These comparisons bring useful context for the analysis of our results. It is clear that the differences between the non-phosphorylated and the mono-phosphorylated proteoforms are much more significant than those between the mono-phosphorylated and the di-phosphorylated proteoforms. This observation is consistent with prior biophysical studies on the effects of phosphorylation on global stability, where the differences in thermodynamic stability between non-phosphorylated and mono-phosphorylated OVA is much greater than between mono-phosphorylated and di-phosphorylated^41^. The addition of the second phospho group increases the magnitude of protection of some residues in the LC-FOX footprint and adds some new sites of protection, but no sites that were protected upon addition of the first phospho group at Ser68 become unprotected upon addition of the second phospho group at Ser344. Specific hot-spots of structural change can be observed adjacent in space, although often distant in sequence. The 138-153 helix shows significant protection at the C-terminal end, as well as protection in multiple loops at the bottom of the structure (as oriented in **Figure 6**) that becomes slightly more extensive upon the addition of the second phosphate. Both partners in the interaction interface between Cys30 and His328 show coordinated protection upon the first phosphorylation event. Most notably, the addition of the second phosphate at Ser344 at the top of the structure induces new protection in the nearby β-strands that are directly adjacent to the coordinated Lys189. The stretch of 348-350 shows no change in protection upon phosphorylation of Ser344, which initially seems contrary to expectations given the stabilizing effect phosphorylation of Ser344 has on this loop. However, the most likely oxidation target in this stretch (Val349) is fully solvent-exposed in the ordered helix, pointing away from the core of the structure. Therefore, stabilization of this helix should not protect Val349 from oxidation. No regions in the protein showed an increase in oxidation upon phosphorylation.

**Figure 6.**
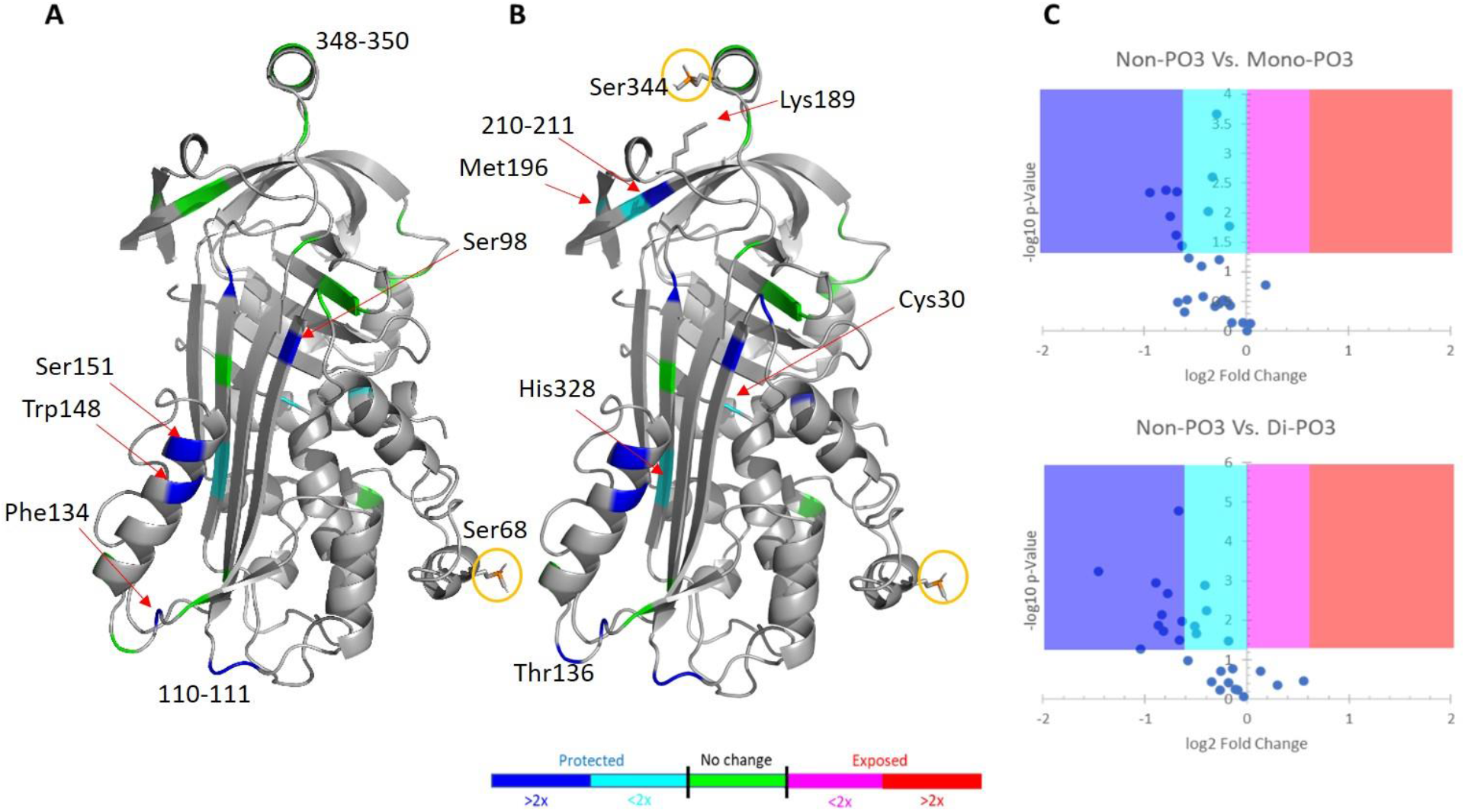
Comparison of residue level LC-FOX data plotted on the structure of OVA. **A)** Change in LC-FOX footprint between non-phosphorylated and mono-phosphorylated OVA. **B)** Change in LC-FOX footprint between non-phosphorylated and di-phosphorylated OVA. Dominant site(s) of phosphorylation are highlighted in a yellow circle. Residue color represents fold change in oxidation, while green residues represent no statistically significant change in oxidation. Grey residues showed no measurable oxidation.

## CONCLUSION

The novel LC-FOX technology described here allowed us to analyze three OVA phosphoproteoforms in an ongoing reaction with alkaline phosphate, and differentiate the impacts of each phosphorylation event on the surface of the protein at the residue level. The data we generated were consistent with both prior biophysical data indicating the role of phosphorylation in stabilization of OVA^41,42,45^, as well as with local MD simulations showing interactions between phosphoSer344 and Lys189 resulting in changes in local dynamics. Regions showing protection are commonly spatially adjacent, even if they are distant in sequence. While these results are consistent with prior data, they offer unique insights into the structural consequences of phosphorylation. The ability to examine changes in protein structure within dynamic systems, now using much higher resolution chromatography than previously achieved for LC-FPOP and a much more robust and convenient illumination source^23^ makes this technology both more impactful and more readily adopted by other laboratories. Demonstration of the lack of radical scavenging by even high concentration NaCl gradients also opens up hydrophobic interaction chromatography (another highly useful option for separation of native protein conformers)^46^ for simple adoption into LC-FOX workflows, without the need for a compensation strategy^47^. Work is ongoing examining the impacts of this technology on other problems of biomedical importance, including complex protein-ligand binding systems, where this technology can help to tease apart structural differences in dynamic binding modes that cannot be purified and characterized separately.

## ASSOCIATED CONTENT

Supplementary Information

## AUTHOR INFORMATION

### Notes

The authors declare the following competing financial interest(s): ZC and JSS disclose a significant interest in GenNext Technologies, Inc., a small company seeking to commercialize technologies for protein higher-order structure, protein-protein interactions, protein-ligand interactions, and PTM analysis.

## ACKNOWLEDGMENTS

The authors would also like to thank GenNext Technology for loaning the FOX photolysis system for this work. This work was supported by a grant from the National Institute of General Medical Sciences (R01GM127267). The authors would like to thank Dr. Sushil K. Mishra of the Glycoscience Center of Research Excellence Computational Chemistry and Bioinformatics Research Core for performing the MD simulation. SEC-MALS and MD simulation analyses were supported by the Glycoscience Center of Research Excellence (P20GM130460).

