## Supplementary Information for "Structural Analysis of Phosphorylation Proteoforms in a Dynamic Heterogeneous System Using Flash Oxidation Coupled In-Line with Ion Exchange Chromatography"

A

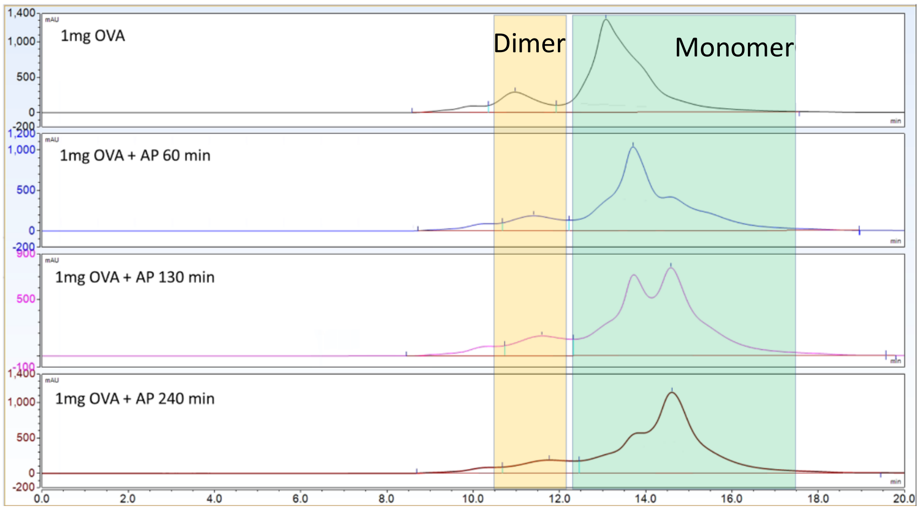


| Name | Dimer Area (mAU*min) | Monomer Area (mAU*min) | Total Area (mAU*min) | % Dimer | % Monomer |
|---|---|---|---|---|---|
| 1mg OVA | 297 | 1770 | 2067 | 85.63 | 14.37 |
| 1mg OVA+AP 60min | 243 | 1576 | 1819 | 86.64 | 13.36 |
| 1mg OVA+AP 130min | 233 | 1593 | 1826 | 87.24 | 12.76 |
| 1mg OVA+AP 240min | 271 | 1881 | 2152 | 87.41 | 12.59 |

B

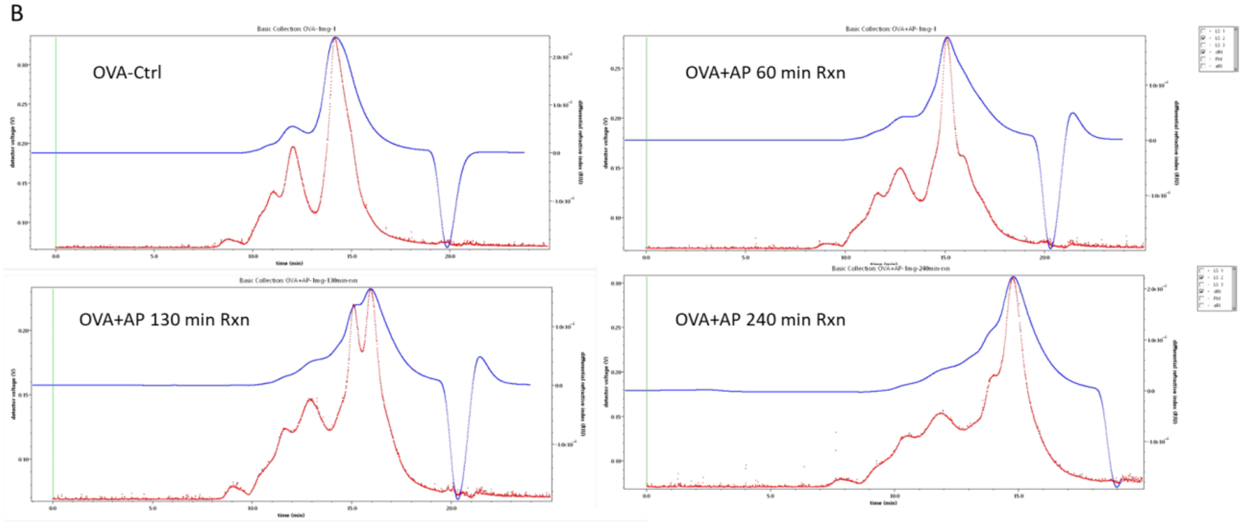


**Figure S1: SEC-MALS analysis of OVA-AP reaction products.** OVA without AP, OVA with AP at time points of 60, 130, and 240 min were analyzed by SEC-MALS. **A)** UV 280 nm absorbance trace from SEC-MALS. The percentage of the OVA monomer and dimer were calculated by the peak area from the UV result, the dimer eluted around 10-12 min (yellow) and the monomer eluted from 12-17 min (green) with at least two conformers of monomer. **B)** The (red) MALS light scattering data and the (blue) refractive index measured for each OVA sample. Early eluting peak was verified as OVA dimer with an estimated molecular weight of 84(±10%) kDa. The late eluting peaks were verified as monomer, with an estimated MW of 42(±5%) kDa and 39(±5%) kDa.

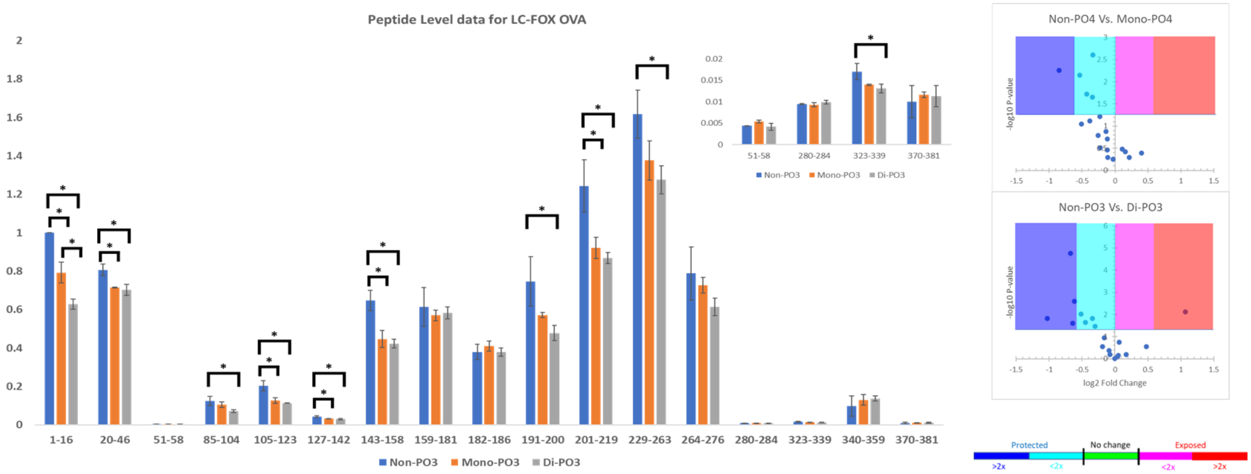


**Figure S2.** Peptide level calculation of three OVA proteoforms. The data were analyzed by one-way ANOVA followed by Tukey post-hoc analysis (α=0.05). Asterisks mark data pairs with statistically significant differences, as indicated by the bracket. **Inset:** A view of peptides with low levels of oxidation, zoomed in on the Y-axis.

**Table 1. LC-MS/MS targeted mass list for OVA residue level analysis.**

| Sequence | Position | m/z | z | t start (min) | t stop (min) |
|---|---|---|---|---|---|
| GSIGAASMEFC[+57.021]FDVFK | 1-16 | 883.401 | 2 | 30 | 32 |
| | 1-16-OX | 891.398 | 2 | 28 | 31 |
| VHHANENIFYC[+57.021]PIAIMSALAMVYLGAK | 20-46 | 759.132 | 4 | 37 | 39 |
| | 20-46-OX | 763.131 | 4 | 35 | 39 |
| TQINKVVR | 51-58 | 479.297 | 2 | 17 | 19 |
| | 51-58-OX | 487.294 | 2 | 15.5 | 18.5 |
| DILNQITKPNDVYSFSLASR | 85-104 | 761.064 | 3 | 27.5 | 29.5 |
| | 85-104-OX | 766.395 | 3 | 27 | 31 |
| LYAEERYPILPEYLQCVK | 105-123 | 762.061 | 3 | 27.5 | 29 |
| | 105-123-OX | 767.392 | 3 | 26.5 | 29.5 |
| GGLEPINFQTAADQAR | 127-142 | 844.423 | 2 | 25 | 27 |
| | 127-142-OX | 852.42 | 2 | 24 | 27 |
| ELINSWVESQTNGIIR | 143-158 | 929.985 | 2 | 28.5 | 30.5 |
| | 143-158-OX | 937.982 | 2 | 26 | 30 |
| NVLQPSSVDSQTAMVLVNAIVFK | 159-181 | 1230.664 | 2 | 33 | 35 |
| | 159-181-OX | 1238.661 | 2 | 31 | 33.5 |
| GLWEK | 182-186 | 316.673 | 2 | 21 | 22 |
| | 182-186-OX | 324.67 | 2 | 19 | 21 |
| AFKDEDTQAMPFR | 187-199 | 519.245 | 3 | 22.5 | 24 |
| | 187-199-OX | 524.576 | 3 | 20 | 22.5 |
| DEDTQAMPFR | 191-200 | 605.264 | 2 | 22.5 | 24.5 |
| | 191-200-OX | 613.261 | 2 | 20 | 22 |
| VTEQESKPVQMMYQIGLFR | 201-219 | 762.053 | 3 | 28.5 | 30.5 |
| | 201-219-OX | 767.384 | 3 | 25.5 | 28 |
| ILELPFASGTMSMLVLLPDEVSGLEQLESIINFEK | 229-263 | 1288.341 | 3 | 38 | 40 |
| | 229-263-OX | 1293.672 | 3 | 36 | 39 |
| LTEWTSSNVMEER | 264-276 | 791.363 | 2 | 23.5 | 25.5 |
| | 264-276-OX | 799.36 | 2 | 21.5 | 25.5 |
| VYLPR | 280-284 | 324.197 | 2 | 19.5 | 21.5 |
| | 280-284-OX | 332.194 | 2 | 18.8 | 22 |
| ISQAVHAAHAEINEAGR | 323-339 | 591.97 | 3 | 18 | 21 |
| | 323-339-OX | 597.302 | 3 | 17 | 22 |
| EVVGSAEAGVDAASVSEEFR | 340-359 | 1004.977 | 2 | 25.5 | 27.5 |
| | 340-359-OX | 1012.974 | 2 | 24 | 28 |
| HIATNAVLFFGR | 370-381 | 673.371 | 2 | 25 | 28 |
| | 370-381-2OX | 689.365 | 2 | 26 | 29 |
